# Growth and Yield Response of Soft White Common Spring Wheat (SWCSW) Varieties under Different Nitrogen Fertilizations and Plant Growth Regulators Applications

**DOI:** 10.1101/2025.09.22.676075

**Authors:** Tajamul Hussain

## Abstract

Application of plant growth regulators (PGRs) is often reported beneficial in wheat to achieve various targets including lodging resistance. However, research evidence on the application of PGRs in Soft White Common Spring Wheat (SWCSW) varieties is limited particularly in Columbia Basin. Therefore, our objectives were to evaluate the effects of PGRs on SWCSW varieties exposed to different nitrogen (N) application levels. Two widely grown SWCSW varieties, Louise and Diva were planted for two growing seasons using split plot design with four replications. Crop growth (stem height and diameter), yield and yield attributes (grain numbers per head, head numbers per fit, grain weight), quality characteristics (test weight, protein content) and lodging score were recorded following the application of PGRs at tillering (GS21-26), stem elongation (GS30-32), and flag leaf emergence (GS37-39) stages combined with high N fertilization. No lodging was observed during first season whereas it was witnessed during second season, and we observed a variable but positive response particularly under application of CC-C, TE, CC-BC and CC-AB. An increase in stem diameter and reduction in stem height were observed under PGR application which indirectly contributed to stem strength and reduced lodging. Although impact of PGRs application on grain numbers, head numbers grain yield and grain weight were non-significant and variable, we observed an improved performance of these attributes such as higher grain yield was observed under application of CC-B and CC-BC. The grain yield of Louise and Diva was similar at low and high N input. These results indicate that application of 168 kg N ha^-1^ is sufficient for an acceptable grain productivity and to gain agronomic returns. As we observed a significant seasonal impact and variations in the performance of SWCSW varieties, future research should consider long term evaluations to evaluate the impacts of PGR’s on soft white common spring wheat varieties.

## 1. Introduction

Wheat is a major cereal and source for human calories and protein and its production and utilization continues to rise (United States Department of Agriculture Foreign Agricultural Service (USDA FAS), 2020). Since the beginning of the farming in Pacific Northwest in late 1800’s, wheat has been the dominant crop in the region (Pan et al., 2017). Soft white common spring wheat (SWCSW) is among the most important cereals in the Columbia Basin of Oregon. Spring wheat productivity is influenced by seasonal climatic changes particularly due to high winds and heat waves. Wheat in Columbia Basin is cropped in rotations with several crops such as potatoes, corn, seed grasses onions and legumes (Qin et al., 2020). High yielding and input responsive wheat varieties contribute to sustainable wheat production. Crop management practices are particularly evolving to combat climate change impacts and cope with seasonal fluctuations in weather patterns. Therefore, it is critical to evaluate the response of widely grown SWCSW varieties to various inputs and management practices such as nitrogen (N) fertilization and application of plant growth regulators (PGR’s).

Nitrogen is a vital nutrient for wheat crops and deficiency of N results in agronomic losses impacting the profitability of wheat cropping systems. Whereas excessive or improper N application has environmental consequences (Ma et al., 2019). Hence, nitrogen nutrition has a direct impact and determination and application of optimal amount of N is critical for sustainable wheat production. Various methods can be utilized to determine optimal N application rate such as soil sampling. Pre-planting soil sampling is essential as it provides the information on soil residual N (Walsh et al., 2022). To determine the N application rate, measuring the plant available N is the root zone is recommended. In this regard, soil samples from the surface layer should also be analyzed for ammonium-N. Following then, the sum of nitrate from soil profile and ammonium-N from the top layer should be accounted for determining N application rate (Horneck et al., 2010).

Lodging in wheat is a serious concern that reduces photosynthesis rate and dry matter production and impacts grain quality in addition to problems in harvest operations (Ma et al., 2012)(Berry, 2019). It is a complicated phenomenon which is defined as the physical bending or breaking of wheat stems. Lodging occurrence can reduce wheat grain yields 8-61% (Berry and Spink, 2012). Lodging is influenced by numerous factors such as wind speed, rainfall, crop management practices and soil conditions (Berry, 2019). Cereals are tend to stem elongation under high N availability and plant densities that also facilitate lodging (Shah et al., 2019). Improved and high yielding varieties required high N input to achieve yield and protein targets. Grain yield of these varieties is often associated with higher grain weight, heavy spikes and number of grains. These factors indirectly contribute to lodging risks as leverage on plant increases compared to root and stem strength. According to Baker et al. (Baker et al., 2014), precipitation, wind and crop management practices impact stem and root stability making prediction of lodging more difficult.

Application of plant growth regulators (PGRs) is considered one of the suitable agronomic practices in addition to adjustment in planting densities and split application of fertilizers (Wu et al., 2019)(Mizuta et al., 2020). Use of PGRs has also contributed to reducing crop heights making cereal more tolerant to lodging risks and improvement in yields in some countries (Savin et al., n.d.). According to VanDerZande (Van Der Zanden, 2012), PGRs are synthetic compounds that simulate plant hormones and interrupt biosynthesis of plant hormones, that alters the plant growth and development. Reducing stem height and increasing stem diameter of wheat and application of PGRs would strengthen the stem ultimately leading to reduced lodging risks. The application of PGRs is more effective when applied at appropriate crop growth and development stage. Split application is also considered useful as it can match the ideal crop stage to induce desired effects. Auxin, gibberellin, cytokinin, ethylene, and abscisic acid are five types of plant hormones that affect plant growth (Anderson et al., 2015). According to Berry et al. (Berry et al., 2004) PGRs particularly those inhibit gibberellin biosynthesis can limit stem elongation and increase stem strength potentially reducing lodging risks under higher N application rates. Role of PGRs in gibberellin biosynthesis inhibition at early stage was observed under application of chlormequat chloride (CC) and mepiquat chloride whereas trinexapac-ethyl (TE) and prohexadione-Ca were affective at lateral stages (Rademacher, 2000). PGRs inhibiting gibberellin such as CC and TE, shorten internodal length and enhance stem strength, thus improving wheat’s stem ability to tolerate high winds. According to Hedden (Hedden and Sponsel, 2015) application of CC in wheat is the largest application to control lodging. Whereas TE was introduced a few decades ago (King et al., 2004). Application rate of TE ranges 841-1009 gha^-1^ whereas ideal timing of application is at the initiation of stem elongation and beginning of flag leaf emergence. Application of TE in Columbia Basin of Oregon has provided benefits to reduce lodging and improve winter wheat yields, however, scientific evidence on its impact on spring wheat is limited. Although the application of PGRs has provided benefits to control lodging, their efficiency is dependent upon application timing, type of PGR, dosage and genetic characteristics of specific wheat varieties. As a result, research work on evaluation of PGR application on SWCSW will provide insights to manage lodging risks in high pressure areas like Columbia Basin.

Keeping in view the importance of above, it is critical to evaluate the response of spring wheat to different N fertilization rates and N fertilization in combination with different PGRs. The guidelines will be useful for regional SWCSW growers and in the similar cropping systems worldwide for better crop management and adjustment in prevailing practices according to soil and climatic conditions. Hence the objectives of our research were to determine i) the impacts of application of PGRs at different crop growth stages on spring wheat productivity and quality through evaluation of crop growth, yield, quality and lodging potential and ii) to evaluate the suitable combination of N fertilization and PGR application through the assessment of their interactions. Findings of the study will ultimately lead to reduced lodging risks and improve grain productivity and grain quality of soft white common spring wheat varieties.

## 2. Materials and Methods

### 2.1. Plant Material

Two widely grown soft white common spring wheat (SWCSW) varieties including Louise and Diva that are popular among farmers in Columbia Basin of Oregon and are regionally well adapted were used in this study. These varieties were chosen due to their resistance against diseases, grain productivity and quality and seed availability. Louise (Reg. No. CV-987, PI 634865) was developed and jointly released in August 2005 by the Agricultural Research Center of Washington State University in collaboration with the experiment stations of the University of Idaho and Oregon State University, and the United States Department of Agriculture-Agricultural Research Service (USDA-ARS). Louise has superior end-use quality, high yield potential, high-temperature adult-plant resistance to local races of stripe rust and partial resistance to the Hessian fly (Kidwell et al., 2006). Diva (Selection No’s. SW02064, S0500302L and Line designation: WA8090) was developed by the Western Wheat Quality Laboratory of USDA-ARS in collaboration with Wheat Quality Program of Department of Crop and Soil Sciences, Washington State University. Diva is a suitable SWCSW variety for commercial production in the areas where the stripe rust and Hessian fly are serious concerns. Diva has an outstanding grain yield potential across a broad range of environments, better resistance against Hessian fly and stripe rust resistance with higher test weight whereas it is shorter in height and greater in flour yield, break flour yield and milling score than Louise. It is well suited for production in the intermediate and high rainfall zones (Morris et al., 2009).

### 2.2. Experimental setup and crop management

Two-year field experiments were conducted at the Hermiston Agricultural Research and Extension Center (45°50′25″ N, 119°17′22″ W, elevation: 140 m) of Oregon State University on an Adkins fine sandy loam (*mesic Fluvaquentic Endoaquepts*) soil. Experimental area belongs to a semi-arid climate with a averaged low and high temperature range of 5 °C-18.9 °C and an annual precipitation of 264 mm (Climate Hermiston–OR, 2020). Randomized complete block design with four replicates was used to layout the experiments during both years. SWCSW varieties including Louise and Diva were planted with planting density of 300 seeds m^−2^ at 1.3 cm planting depth on April 2 and April 6 during first (2020) and second (2021) season, respectively. Each variety was grown with a cone planter holding 9 rows in 2020 and 7 rows in 2021 with a 2-m wide pass. There were 8 treatments in the experiments consisting of normal N rate (N1), high N rate (N2), as well as five plant growth regulators treatments. All the PGR treatments were based on the high N scenario including test PGR (CC) applied at tillering, 2-node, and flag leaf stage alone or split, representing as CC-A, CC-B, and CC-C, CC-AB, CC-BC, and Palisade (TE). Prior to planting the crop, fertilizer consisted of 168 kg N ha^−1^ and 22 kg S ha^−1^ was broadcast-applied on soil surface (normal N rate, N1) which was consistent with farmers’ practices in the region. During tiller stage, additional 112 kg N ha^−1^ was applied to most treatments to achieve the higher N scenario (N2) which was used to promote the occurrence of lodging and to thoroughly test the efficacy of the PGR products. Central pivot irrigation system was used to apply irrigation that was scheduled during April to June in each season.

### 2.3. Application of Plant Growth Regulators (PGRs)

The PGR products included Palisade (a.i. 12% TE) and Adjust (Pending Registration; a.i. 55% CC). The PGR were applied at three BBCH plant growth stages including tillering (GS21-26), stem elongation (GS30-32), and flag leaf (GS37-39) (Table 1). Application of PGR and related growth stages are summarized in Table 1. Control treatments for low and high N application rates did not receive any PGR applications while PGR combinations were applied with high N application rates at the specified stages. Backpack sprayer was used to apply PGR on crop foliage.

**Table 1.** Treatment details for the two-year field trials in Oregon.

| Treatment | Treatment Codes | Nitrogen Application | Application of PGR's at Different Crop Growth Stages |  |  |
| --- | --- | --- | --- | --- | --- |
|  |  |  | GS21-26<br>(Tillering) | GS30-32<br>(Stem Elongation) | GS37-39<br>(Flag Leaf Emergence) |
| 1 | N1 | Low N rate | 0 | 0 | 0 |
| 2 | N2 | High N rate | 0 | 0 | 0 |
| 3 | CC-A | High N rate | Test PGR 25 fl oz | 0 | 0 |
| 4 | CC-B | High N rate | 0 | Test PGR 25 fl oz | 0 |
| 5 | CC-C | High N rate | 0 | 0 | Test PGR 25 fl oz |
| 6 | TE | High N rate | Palisade 14.4 fl oz | 0 | 0 |
| 7 | CC-BC | High N rate | 0 | Test PGR 14 fl oz | Test PGR 14 fl oz |
| 8 | CC-AB | High N rate | Test PGR 11 fl oz | Test PGR 14 fl oz | 0 |

### 2.4. Data collection

Stem height (cm) and stem diameter (mm) were recorded twice in both growing seasons. First at the water ripe when first grains reached half of their maximum size (GS 71) on 25 May and on 3 June, and at physiological maturity (GS92) on 25–26 June and on 15–17 July for first and second season, respectively. Stem height was recorded from 10 plants in each plot from the base of the stem to the top of head excluding the awn. Stem diameter was measured 1” above the ground on main stem using automated caliper and total of 10 plants were used to obtain data on stem diameter. Harvesting was performed using small-plot harvester and experimental plots were harvested on 7 July and 30 July for the first and second season, respectively. Plants from one central row in a length of 0.5 m in each plot were cut above ground prior to a week before harvesting. Head numbers per meter, thousand grain weight (g), grain moisture (%), grain yield (Mt ha^−1^), protein content (%) and test weight (kg L^−1^) were recorded. The grain number per head was counted from 10 randomly selected heads. The weight of a thousand grain was obtained from two random samples of 50 grains from each treatment sample and was converted to 1000 grain weight (g). Based on visual field observations, the lodging score was recorded. The lodging score was based on the scale of 1-9 with serious lodging at 9 and no lodging occurrence at 1.

### 2.5. Data analysis

Data on crop growth and yield parameters was analyzed using Statistix 8.1 package (Analytical software, Tallahassee, USA) (Duangpan et al., 2022) and analysis of variance (ANOVA) was computed for stem height at water ripe, stem height at harvest maturity, stem diameter at water ripe, stem diameter at harvest maturity, grain numbers per head, head numbers per fit, 1000 grain weight, grain yield, protein contents, test weight and lodging score of Louice and Diva soft white common spring wheat (SWCSW) varieties in response to applied treatments and their interactions over two years. Least significant difference (LSD) was used for mean comparisons and a *p*-value less than 0.05 was considered as significant. Pearson’s correlation coefficients were computed for wheat parameters separately for further insights for nitrogen fertilization and for nitrogen in combination with PGR’s application treatments separately using Corrplot package of R program.

## 3. Results and Discussion

### 3.1. Seasonal Effect and Crop Performance

Seasonal effects were significant in statistical analysis possibly due to prevailing climatic conditions (Figure 1) and varietal performance differed over the years (Table 2). Mean daily minimum and maximum temperatures ranged from 27 to 68 and 55 to 98 °F during the first and 25–79 and 56–112 °F during the second growing year, respectively. June and July were the hottest months where the highest minimum and maximum temperatures were observed. Total rainfall during the first and second growing year was 2.17″ and 0.62″ respectively whereas average windspeed was similar over the years with 6 mph. Maximum windspeed was observed during April and June in the first (at 16 mph) and during June in second growing year (at 15 mph), respectively. Higher mean minimum and maximum temperatures and wind speed during the months of June and July when the crop was at lateral and maturity stages possibly impacted wheat causing lodging in the second season. Higher temperatures affect wheat crop in various ways such as causing a reduction in stem strength and reduced root anchoring (CROOK and ENNOS, 1993; Berry et al., 2000) in addition to complex interactions of higher temperatures with soil moisture that play a role in lodging vulnerability.

**Table 2.** F–values and significance obtained through analysis of variance of agronomic attributes including stem height at water ripe (SHWR: GS71) and stem height at harvest maturity (SHHM:GS92), stem diameter at water ripe (SDA: GS71) and stem diameter at harvest maturity (SDB: GS92), grain numbers per head (GN), head numbers per fit (HN), 1000 grain weight (GW), grain yield (GY), protein contents (PC), test weight (TW) and lodging score (LS) of Louise and Diva soft white common spring wheat (SWCSW) varieties.

| Traits | Variety (V) | Treatment (T) | Year (Y) | V×T | V×Y | T×Y | V×T×Y | CV |
| --- | --- | --- | --- | --- | --- | --- | --- | --- |
| SHWR | 31.93<sup>***</sup> | 2.39<sup>*</sup> | 156.80<sup>***</sup> | 0.34<sup>ns</sup> | 5.25<sup>*</sup> | 1.47<sup>ns</sup> | 0.65<sup>ns</sup> | 3.94 |
| SHHM | 8.54<sup>**</sup> | 2.70<sup>*</sup> | 271.26<sup>***</sup> | 0.80<sup>ns</sup> | 10.44<sup>**</sup> | 3.20<sup>**</sup> | 0.67<sup>ns</sup> | 4.55 |
| SDA | 1.08<sup>ns</sup> | 1.3<sup>ns</sup> | 302.32<sup>***</sup> | 1.61<sup>ns</sup> | 2.16<sup>ns</sup> | 2.12<sup>*</sup> | 0.51<sup>ns</sup> | 5.51 |
| SDB | 0.17<sup>ns</sup> | 0.67<sup>ns</sup> | 799.61<sup>***</sup> | 1.89<sup>ns</sup> | 1.09<sup>ns</sup> | 0.87<sup>ns</sup> | 2.14<sup>*</sup> | 8.06 |
| GN | 0.27<sup>ns</sup> | 0.53<sup>ns</sup> | 4.01<sup>*</sup> | 1.01<sup>ns</sup> | 0.05<sup>ns</sup> | 0.76<sup>ns</sup> | 0.87<sup>ns</sup> | 15.89 |
| HN | 2.77<sup>ns</sup> | 1.24<sup>ns</sup> | 61.24<sup>***</sup> | 1.49<sup>ns</sup> | 0.38<sup>ns</sup> | 0.70<sup>ns</sup> | 0.79<sup>ns</sup> | 26.75 |
| GY | 22.97<sup>***</sup> | 0.77<sup>ns</sup> | 139.69<sup>***</sup> | 0.68<sup>ns</sup> | 8.84<sup>***</sup> | 0.34<sup>ns</sup> | 0.80<sup>ns</sup> | 18.91 |
| GW | 23.48<sup>***</sup> | 0.97<sup>ns</sup> | 121.56<sup>***</sup> | 0.63<sup>ns</sup> | 20.76<sup>***</sup> | 0.34<sup>ns</sup> | 0.73<sup>ns</sup> | 18.65 |
| PC | 0.98<sup>ns</sup> | 4.81<sup>***</sup> | 108.97<sup>***</sup> | 0.84<sup>ns</sup> | 8.77<sup>**</sup> | 0.79<sup>ns</sup> | 0.38<sup>ns</sup> | 3.27 |
| TW | 10.90<sup>***</sup> | 1.00<sup>ns</sup> | 11.54<sup>***</sup> | 0.06<sup>ns</sup> | 6.52<sup>***</sup> | 1.80<sup>ns</sup> | 0.09<sup>ns</sup> | 13.99 |
| LS | 5.42<sup>*</sup> | 0.67<sup>ns</sup> | 572.98<sup>***</sup> | 0.37<sup>ns</sup> | 5.42<sup>*</sup> | 0.67<sup>ns</sup> | 0.37<sup>ns</sup> | 35.12 |

\*\*\*: $p < 0.001$ , \*\*: $p < 0.01$ , \*: $p < 0.05$ , ns: non-significant difference at $p \geq 0.05$ .

**Figure 1.**
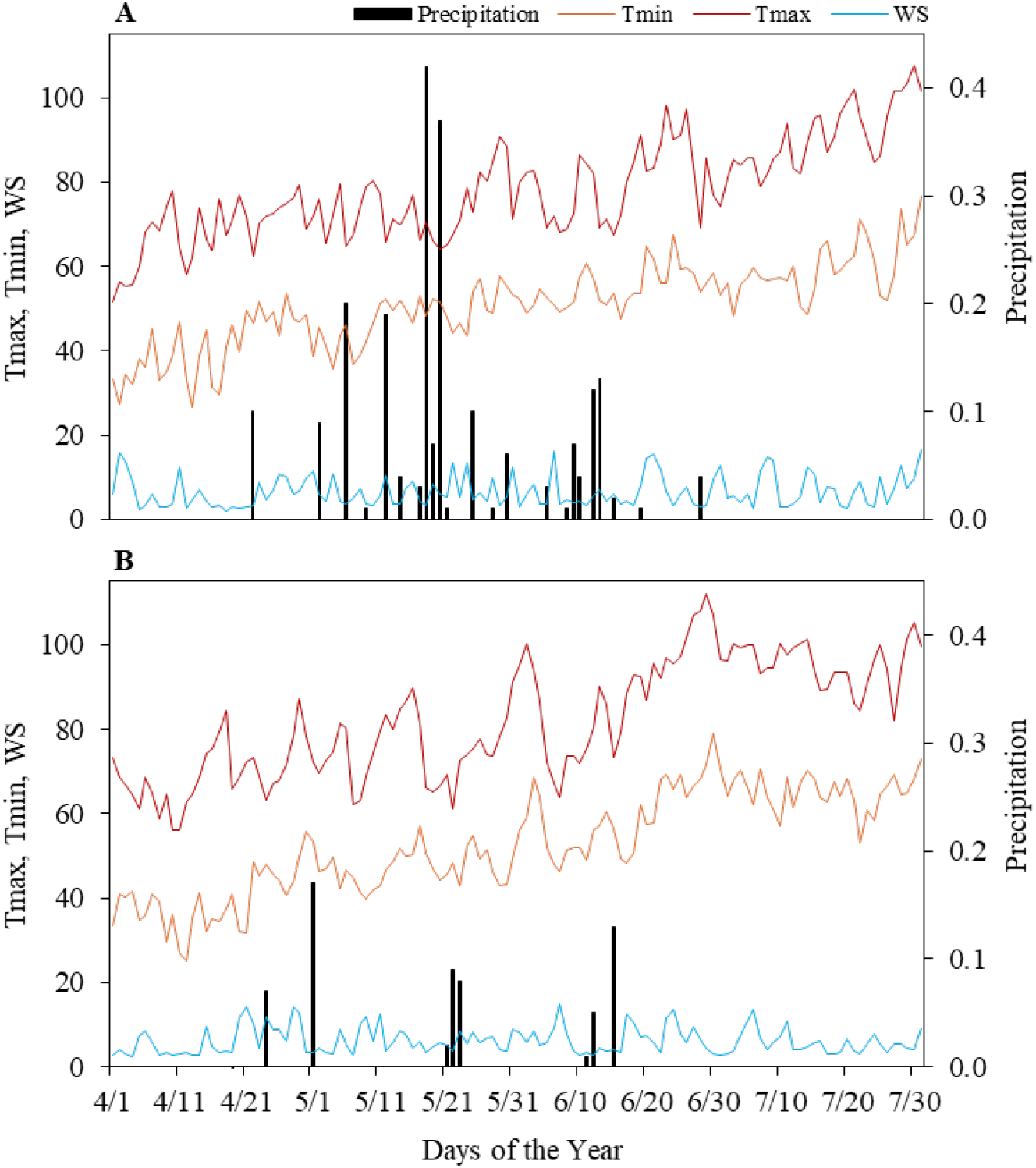
Mean daily maximum (Tmax) and minimum (Tmin) temperatures (°F), daily precipitation (in day^−1^) and daily average wind speed (WS) (mph) during the experimental period (April-July) of the first season: 2020 (A) and second season: 2021 (B). (Data source: Hermiston, Oregon AgriMet Weather Station (HRMO: 45.82111 N; -119.52138 W): Hermiston Agricultural Research and Extension Center (HAREC). AgriMet Cooperative Agricultural Weather Network: Columbia-Pacific Northwest Region, Oregon, United States).

The statistical analysis indicated that applied treatments (T), under the effects of planted varieties (V) and growing years (Y) significantly influenced the stem height at water ripe (SHWR: GS71) and stem height at harvest maturity (SHHM:GS92), 1000 grain weight, grain yield, protein contents, test weight and lodging score (Table 2). There were no significant differences observed for the two-way interactions of variety and growing seasons. Under the influence of interactions between applied treatments and growing years (T×Y), stem height at harvest maturity (SHHM:GS92) and stem diameter at water ripe (SDA: GS71) were only significantly impacted among studied attributes. Stem diameter at harvest maturity (SDB: GS92) among other parameters was only influenced in three-way interaction of variety, applied treatments and growing years (V×T×Y) (Table 2).

### 3.2. Stem Height

Stem height at water ripe and harvest maturity was significantly affected for both varieties. Change in stem height at water ripe was decreased for Louise under PGR’s application that ranged 1-8%. Maximum decline was observed under the application of CC-C by 8% whereas no change was observed in application of TE during first season (Figure 2A). In the second season, stem height at water ripe was decreased only by 1-2% where no change was observed under N2, CC-B and CC-C (Figure 2B). In contrast, it was slightly higher by 1% under CC-A. Diva indicated variable response during both seasons and an increase in stem height at water ripe was observed under N2, CC-A and TE by 1-2% whereas decline in height was observed by 2-5% during first season (Figure 2A). In the second season, stem height at water ripe was increased under CC-A, CC-B and CC-BC ranging 1-3% and was decreased under CC-C and CC-AB ranging 2-3% whereas no difference was observed under N2 and TE (Figure 2B).

**Figure 2.**
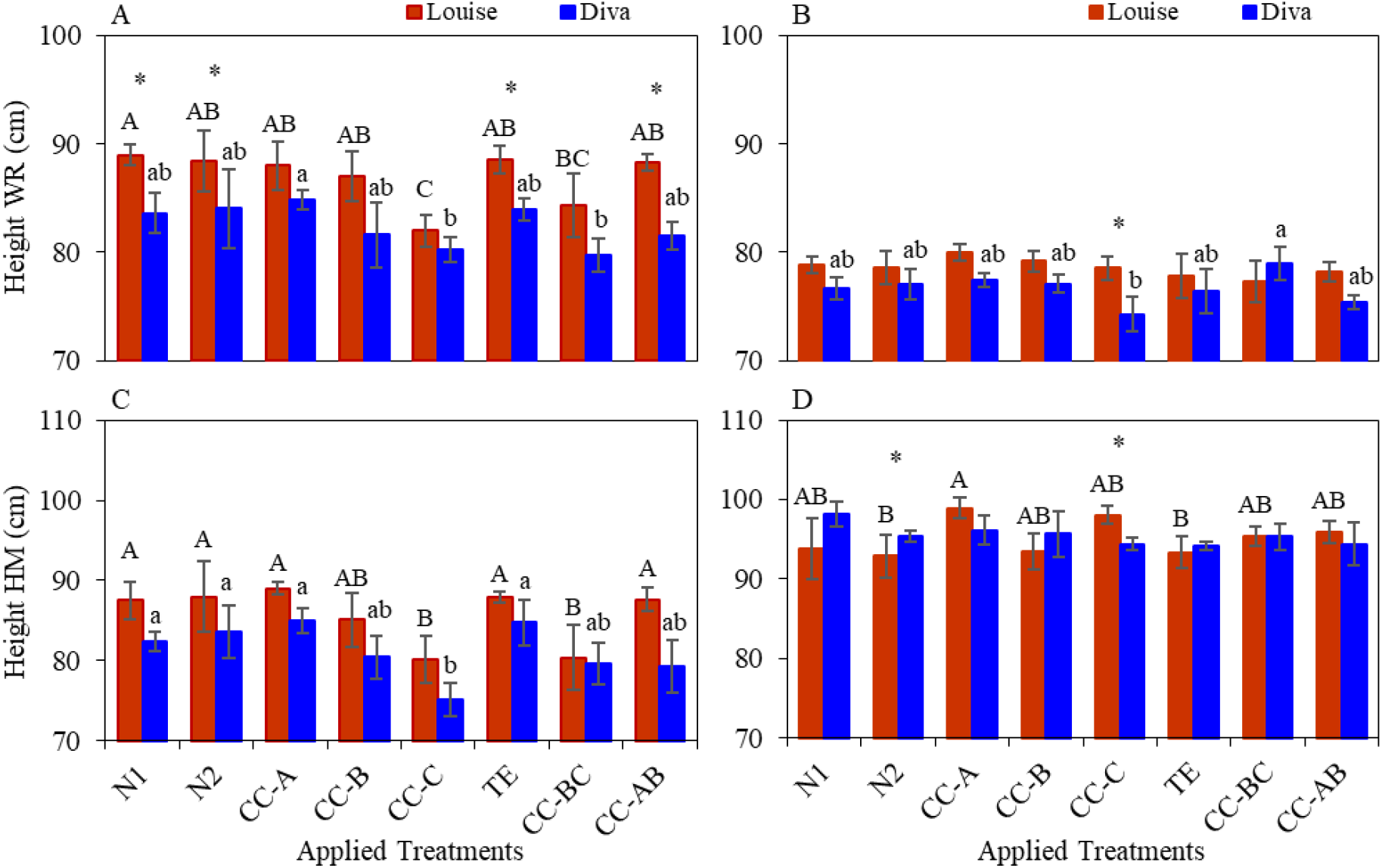
Stem height (cm) at water ripe (GS71: A, B) and harvest mature (GS95: C, D) for the two soft white common spring wheat (SWCSW) varieties among nitrogen and PGR’s treatment combinations during 2020 (A, C) and 2021 (B, D) in Adkins fine sandy loam soil in Oregon. N1 = standard N rate; N2 = high-N rate without PGR application; CC-A = application of CC at GS2126; CC-B = application of CC at GS30-32; CC-C = application of CC at GS37-39; TE = application of TE at GS21-26; CC-BC = application of CC at GS30-32 and GS37-39; CC-AB = application of CC at GS21-26 and GS30-32. Vertical bars indicate ± standard errors for means of four experimental replicates. Uppercase letters indicate the significant (*p* < 0.05) differences among applied treatments for SWCSW Louise. Lowercase letters indicate the significant (*p* < 0.05) differences among applied treatments for SWCSW Diva. Significant differences between two SWCSW varieties at specific applied treatments are indicated by the centered stars above each pair of the graph bars. Bars without the letters or stars indicate no statistically significant difference.

Stem height at harvest maturity was variable for both varieties in both seasons except Diva in the second seasons where it was decreased under all treatments compared to low nitrogen application rate (N1). Stem height at harvest maturity decreased ranging 3-8% under CC-B, CC-C and CC-AB, only increased under CC-A by 2% whereas no effect was observed under N2, TE and CC-AB for Louise in first season (Figure 2C). In the second season, stem height at harvest maturity decreased only 1% where no change was observed under CC-B (Figure 2D). Diva indicated a decline ranging 2-9% under CC-B, CC-C and CC-AB during the first season (Figure 2C). In the second season, it was decreased under all treatments ranging 2-4 (Figure 2D).

Our results indicated that lowest stem height at water ripe was observed under application of CC-C and CC-BC for both varieties in the first and under TE, CC-BC for Louise and under CC-C and CC-AB for Diva in the second season respectively. Considering the stem height at harvest maturity, application of CC-C, CC-BC and CC-AB and CC-B and TE resulted in lowest height in the first and second season respectively. Compared to N1 and N2, application of PGR’s including CC-C, CC-BC and TE might be good strategy to limit the stem height. Although no single PGR can be defined as the best performing in reducing stem height, still application of these PGR’s indicated impact of SWCSW varieties in our study. Several studies have reported the positive impact of application of PGR’s in reducing stem height of wheat varieties (Spolidorio and Lollato, 2019)(Zhang et al., 2017).

### 3.3. Grain Yield

Grain yield of both varieties was significantly different over the seasons (Table 2). However, no significant impact was observed under applied treatments. Grain yield of Louise was significantly higher that Diva under all treatments in both seasons (Figure 3). Highest yield for Louise in the first season (3.9 MT ha^-1^) was observed under higher N application (CC-BC) whereas in the second season (3.4 MT ha^-1^), it was observed under higher N application rate (CC-B) (Figure 3A). Conversely, the lowest yield for Louise in the first season (3.5 MT ha^-1^) was observed under higher N application (TE) whereas in the second season (2.6 MT ha^-1^), it was observed under higher N application rate (N2) (Figure 3A). The highest yield for Diva in the first season (3.7 MT ha^-1^) was observed under higher N application (CC-B) whereas in the second season (3.9 MT ha^-1^) it was observed under low N application (N1) (Figure 3B). However, the lowest yield for Diva in the first season (3.3 MT ha^-1^) was observed under higher N application (CC-AB) whereas in the second season (1.8 MT ha^-1^), it was observed under higher N application rate (CC-C) (Figure 3B).

**Figure 3.**
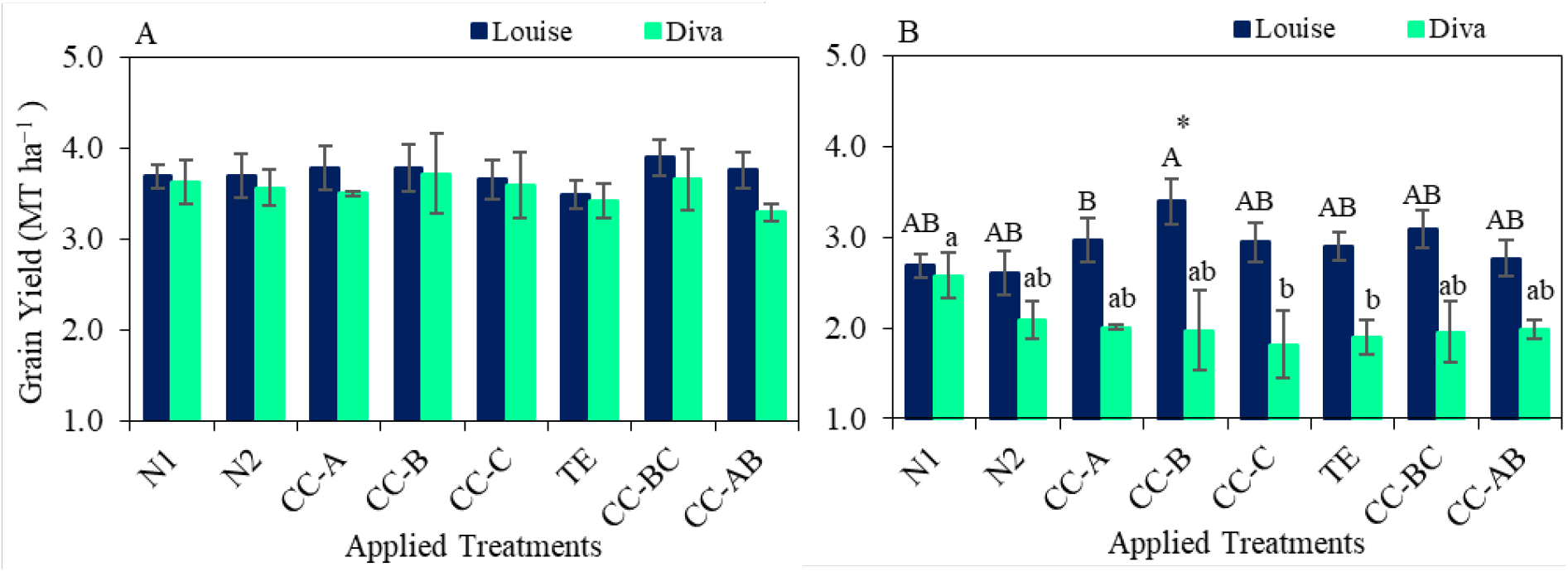
Impact of nitrogen and PGR’s treatment combinations on grain yield of two soft white common spring wheat (SWCSW) varieties during 2020 (A) and 2021 (B) in Adkins fine sandy loam soil in Oregon. N1 = standard N rate; N2 = high-N rate without PGR application; CC-A = application of CC at GS2126; CC-B = application of CC at GS30-32; CC-C = application of CC at GS37-39; TE = application of TE at GS21-26; CC-BC = application of CC at GS30-32 and GS37-39; CC-AB = application of CC at GS21-26 and GS30-32. Vertical bars indicate ± standard errors for means of four experimental replicates. Uppercase letters indicate the differences among applied treatments for SWCSW Louise. Lowercase letters indicate the differences among applied treatments for SWCSW Diva. Significant differences between two SWCSW varieties at specific applied treatments are indicated by the centered stars above each pair of the graph bars.

The grain yield of Louise and Diva was similar under N1 and N2. These results indicate that low N supply (168 kg N ha^-1^) is sufficient for an acceptable grain productivity and reducing N input will result in agronomic and economic benefits. Both varieties were unique in their yield response indicating that they acquired distinct traits that could be explored for further varietal improvements. Grain yield was not significantly different under applied treatments; however, application of CC-BC and CC-B for Louise and CC-BC for Diva were among high yielding treatments in the first and second season respectively. Increase in wheat grain yield is documented in previous studies. For example, Peake et al. (Peake et al., 2020) reported an increase in grain yield of spring wheat varieties particularly under higher N supply. We also witnessed a decline in grain yield of Lousie and Diva under some treatments, such as in the first season application of TE and CC-AB results in lowest yields for Louse and Diva, respectively. However, this reduction was not significant as reported earlier. Previously conducted studies also reported that grain yield of wheat varieties decreased under application of PGR’s (Peake et al., 2020)(Zhang et al., 2017).

### 3.4. Grain Weight

One thousand grain weight of both varieties was also significantly different over the seasons (Table 2). However, no significant impact was observed under applied treatments. In general, the grain weight of Louise was significantly higher than Diva, however under N2, CC-C and TE, the grain weight of Diva was higher than that of Louise during the first season. The highest grain weight for Louise was observed at N1 (37 g) and at N2 (30 in the first and second season, respectively (Figure 4). Conversely, the lowest grain weight for Louise in the first season (30 g) was observed under CC-C (Figure 4A) whereas in the second season (25 g), it was observed under CC-B (Figure 4B). The grain weight of Louise decreased under all treatments except CC-AB compared to N1 in the first season whereas it was variable under different treatments in second season. The grain weight of Diva variable under different treatments in the first season decreased under all treatments compared to N1 in second season. The highest grain weight for Diva was observed at N2 (35 g) and at N1 (23 g) in the first and second season, respectively (Figure 4). Conversely, the lowest grain weight for Diva in the first season (30 g) was observed under CC-AB (Figure 4A) whereas in the second season (25 g), it was observed under application of TE (Figure 4B).

**Figure 4.**
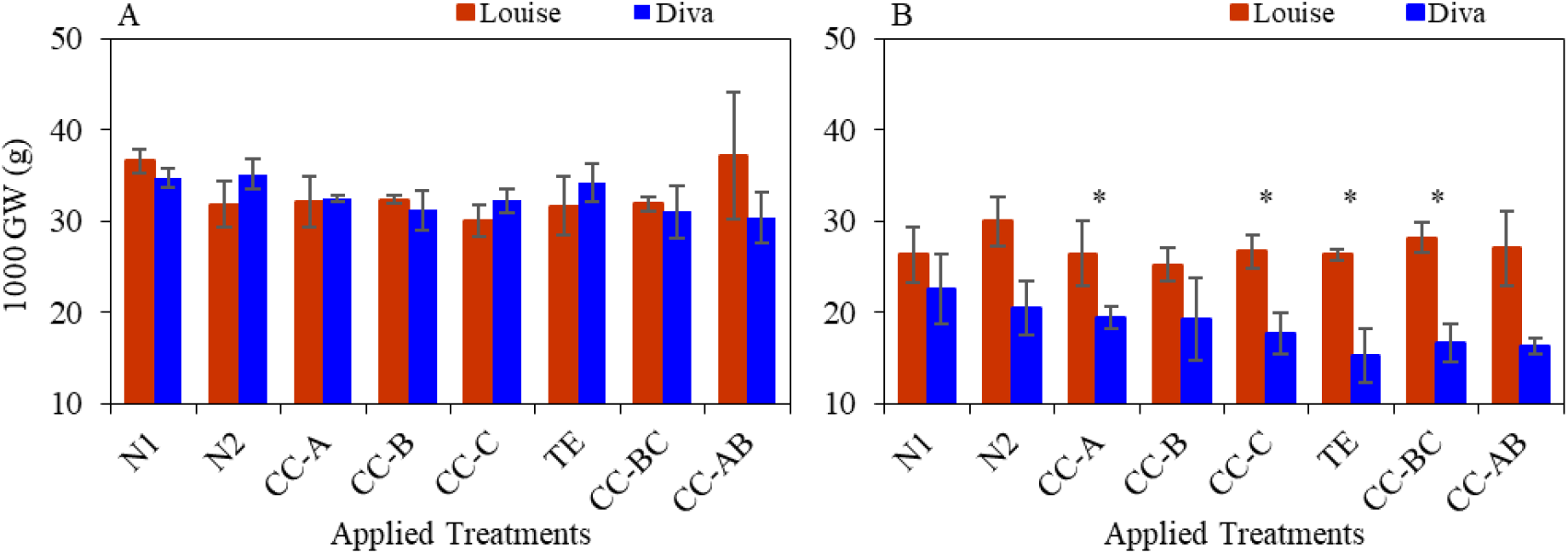
Impact of nitrogen and PGR’s treatment combinations on 1000 grain weight (GW) of two soft white common spring wheat (SWCSW) varieties during 2020 (A) and 2021 (B) in Adkins fine sandy loam soil in Oregon. N1 = standard N rate; N2 = high-N rate without PGR application; CC-A = application of CC at GS2126; CC-B = application of CC at GS30-32; CC-C = application of CC at GS37-39; TE = application of TE at GS21-26; CC-BC = application of CC at GS30-32 and GS37-39; CC-AB = application of CC at GS21-26 and GS30-32. Vertical bars indicate ± standard errors for means of four experimental replicates. Significant differences between two SWCSW varieties at specific applied treatments are indicated by the centered stars above each pair of the graph bars. Bars without the letters or stars indicate no statistically significant difference.

In general, no significant effect of PGR’s was observed in our study. However, we observed a decline in grain weight of Louise in the first and of Diva in the second season. Rajala and Sainio, (Rajala, 2001) reported reduction in grain weight which ultimately resulted in lower grain yields. In our study, we also witnessed similar impact of some PGR’s where lower grain yields had lower grain weights. Grain yield is positively correlated with grain weight as indicated in correlation study (Figure 8,9). Grain weight of Louise was decreased under application of PGR’s particularly in first seasons and correlation analysis indicated that intensity of correlation between grain weight and grain yield of Louise was also decreased under application of PGR’s (Figure 8 A,B).

### 3.4. Protein Content

Both varieties were similar in terms of protein contents and no significant varietal difference was observed (Table 2). However, application of various treatments significantly impacted protein contents of both varieties, and the seasonal effect was also significant (Table 2). The highest protein contents for Louise were 16% and 15% which were observed under TE in both seasons, (Figure 5). Conversely, the lowest protein content of 14% for Louise was observed under N1 in both seasons (Figure 5). The protein contents of Diva were variable under applied treatments and over the seasons and ranged between 14-16 %. Application of various treatments slightly increased protein contents in both seasons. The lowest protein contents (14%) for Diva were observed under N1 in both seasons (Figure 5). Our results indicated that protein contents were higher under higher N application and application of PGR’s compared to low N application. This is usually expected that high N availability results in higher protein and gluten content in grains. Pleskachiov et al. (Pleskachiov et al., 2022) and Yang et al. (Yang et al., 2018) reported similar results that increase in N supply increased the protein contents in wheat.

**Figure 5.**
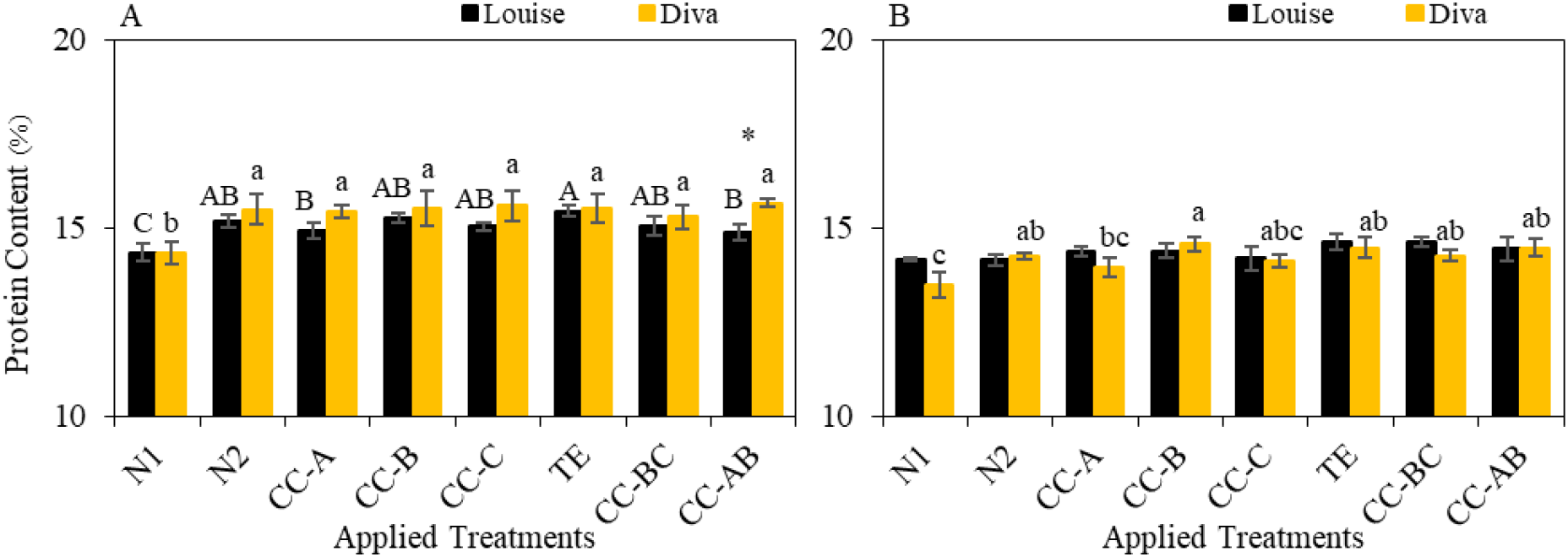
Impact of nitrogen and PGR’s treatment combinations on protein contents of two soft white common spring wheat (SWCSW) varieties during 2020 (A) and 2021 (B) in Adkins fine sandy loam soil in Oregon. N1 = standard N rate; N2 = high-N rate without PGR application; CC-A = application of CC at GS2126; CC-B = application of CC at GS30-32; CC-C = application of CC at GS37-39; TE = application of TE at GS21-26; CC-BC = application of CC at GS30-32 and GS37-39; CC-AB = application of CC at GS21-26 and GS30-32. Vertical bars indicate ± standard errors for means of four experimental replicates. Uppercase letters indicate the significant (*p* < 0.05) differences among applied treatments for SWCSW Louise. Lowercase letters indicate the significant (*p* < 0.05) differences among applied treatments for SWCSW Diva. Differences between two SWCSW varieties at specific applied treatments are indicated by the centered stars above each pair of the graph bars.

### 3.5. Test Weight

Test weight significantly different for both varieties (Table 2). However, application of various treatments indicated no significant impact on test weight of both varieties. In general, the test weight of Louise was higher under all treatments than that of Diva in both seasons (Figure 6). Test weight for Louise ranged 0.68-0.71 kg L^-1^ in the first and 0.54-0.72 kg L^-1^ in the second respectively. The highest test weight for Louise was observed under N1 (0.71 kg L^-1^) in the first and under N2 and CC-C (0.71 kg L^-1^) in the second season, respectively. Test weight for Diva ranged 0.67-0.71 kg L^-1^ in the first and 0.47-0.62 kg L^-1^ in the second respectively. The highest test weight for Diva was observed under N1 (0.71 kg L^-1^) in the first and under N2, CC-C and CC-BC with value of 0.71 kg L^-1^ in the second season, respectively (Figure 6). Both varieties were unique in their test weight. In the first season the test weight of both varieties was similar whereas in the second season it was lower for Diva compared to Louise. In the second season, we observed increase in test weight of Louise and Diva under PGR’s application. Similar findings were also reported by Isaychev et al. (Isaychev et al., 2020) where application of Crezacin and Energia increased the test of spring wheat variety Zemlyachka.

**Figure 6.**
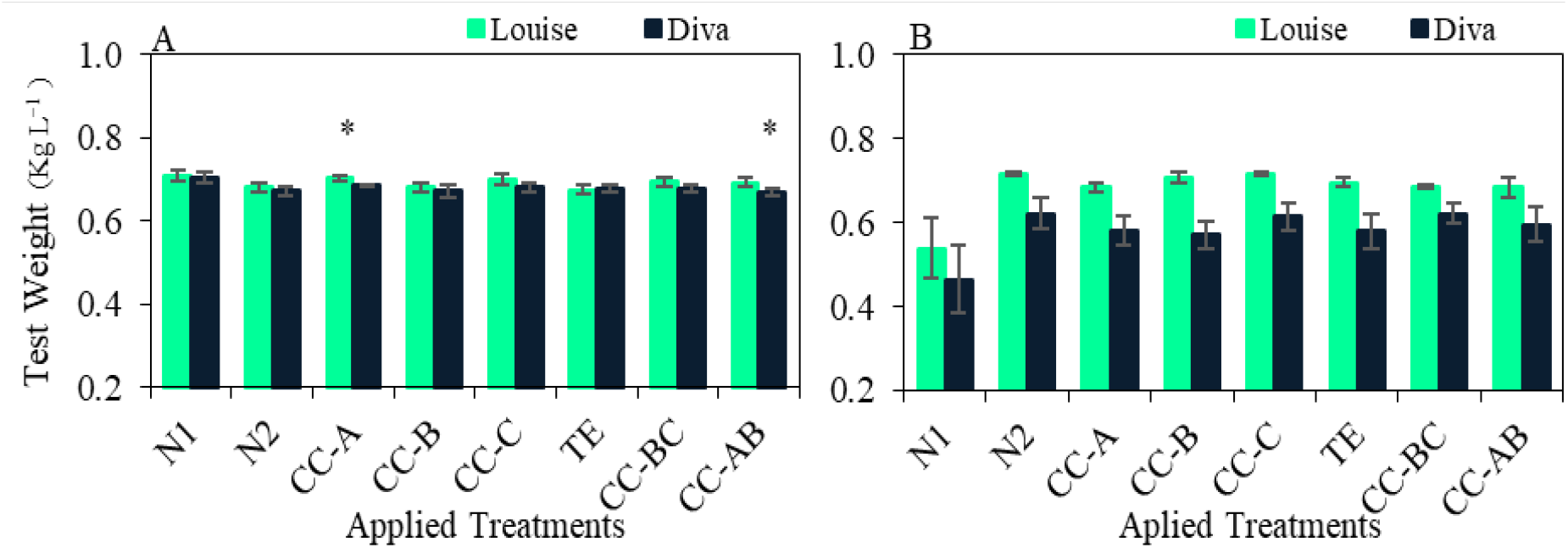
Impact of nitrogen and PGR’s treatment combinations on test weight of two soft white common spring wheat (SWCSW) varieties during 2020 (A) and 2021 (B) in Adkins fine sandy loam soil in Oregon. N1 = standard N rate; N2 = high-N rate without PGR application; CC-A = application of CC at GS2126; CC-B = application of CC at GS30-32; CC-C = application of CC at GS37-39; TE = application of TE at GS21-26; CC-BC = application of CC at GS30-32 and GS37-39; CC-AB = application of CC at GS21-26 and GS30-32. Vertical bars indicate ± standard errors for means of four experimental replicates. Differences between two SWCSW varieties at specific applied treatments are indicated by the centered stars above each pair of the graph bars. Bars without stars indicate no statistically significant difference.

### 3.6. Lodging

Lodging score was significantly different between Louise and Diva whereas application of different treatments did not indicate a direct impact on lodging score (Table 2). During the first season, lodging was not observed for both varieties (LS=1) possibly due to overall shorter stem height in this season. However, lodging occurred in the second season (Figure 7). The lodging score ranged 7-8 for Louise and 5-7 for Diva in the second season (Figure 7). Application of PGR’s resulted in lower lodging score indicating positive impact on lodging resistance. Although the response of various PGR’s in terms of lodging score was variable for both varieties, it can be summarized that application of PGR’s resulted in better plant resistance to lodging. Lodging scores were negatively correlated with stem diameter which indicates that the thicker stems had better resistance against lodging. Lodging can be influenced by several factor such as stem thickness or diameter, stem height, grain weight and overall plant health and climatic conditions. Mangin et al. (Mangin and Mangin, 2022) reported that application of PGR’s impacted on crop canopy thus reducing lodging risks in spring wheat in Canada.

**Figure 7.**
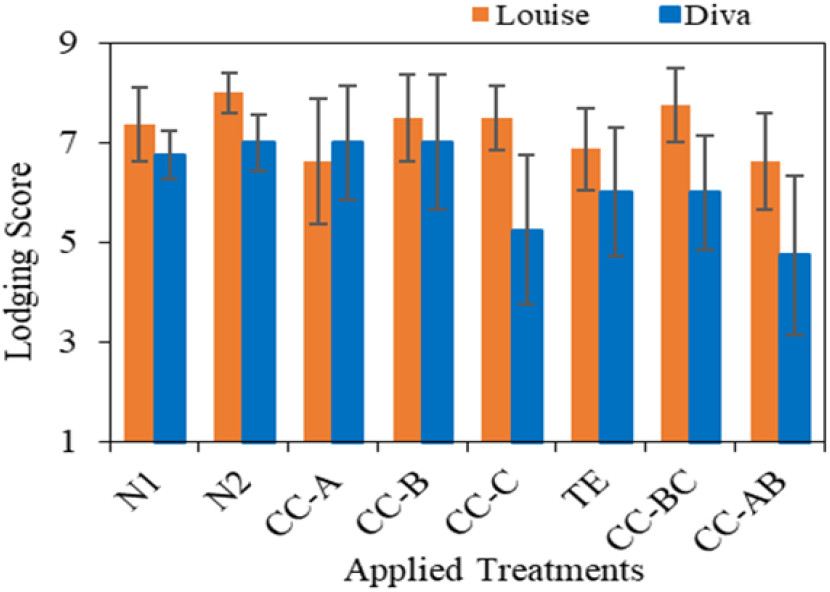
Impact of nitrogen and PGR’s treatment combinations on test weight of two soft white common spring wheat (SWCSW) varieties during 2021 in Adkins fine sandy loam soil in Oregon. N1 = standard N rate; N2 = high-N rate without PGR application; CC-A = application of CC at GS2126; CC-B = application of CC at GS30-32; CC-C = application of CC at GS37-39; TE = application of TE at GS21-26; CC-BC = application of CC at GS30-32 and GS37-39; CC-AB = application of CC at GS21-26 and GS30-32. Vertical bars indicate ± standard errors for means of four experimental replicates.

### 3.7. Correlation Assessment

Correlation analysis indicated the association among the attributes of Louise and Diva differed under N fertilization and under application of PGR’s. Considering N fertilization for Lousie (Figure 8), grain weight was positively correlated with grain numbers (0.39), stem diameter at water ripe (0.52), stem diameter at harvest maturity (0.42), stem height at water ripe (0.41) and grain yield (0.49). Lodging score was negatively correlated with grain numbers (-0.31), grain weight (-0.51), stem diameter at water ripe (-0.92), stem diameter at harvest maturity (-0.87), stem height at water ripe (-0.84) and protein contents (-0.57) whereas it was positively correlated with stem height at harvest maturity (0.50), head numbers (0.74) and test weight (0.54). Stem diameter at water ripe was positively correlated with grain yield (0.76), stem diameter at harvest maturity (0.87), stem height at water ripe (0.80) and protein contents (0.58) whereas it was negatively correlated with, head numbers (-0.76) and test weight (-0.60). Grain yield was positively correlated with grain weight (0.49). Test weight indicated a negative association with stem height at water ripe (-0.63). Considering the effect of PGR’s, degree of association among Louise parameters generally decreased in negative correlations and increased for positive correlations (Figure 8). Association of lodging score was decreased with stem diameter at water ripe (-0.68), stem diameter at harvest maturity (-0.83), stem height at water ripe (-0.68) whereas positive association of lodging score was increased with stem height at harvest maturity (0.73) and decreased with head numbers (0.42). A similar pattern was also observed among other attributes such as stem diameter at water ripe, stem diameter at harvest maturity, stem height at water ripe, head numbers, protein contents and test weight (Figure 8).

**Figure 8.**
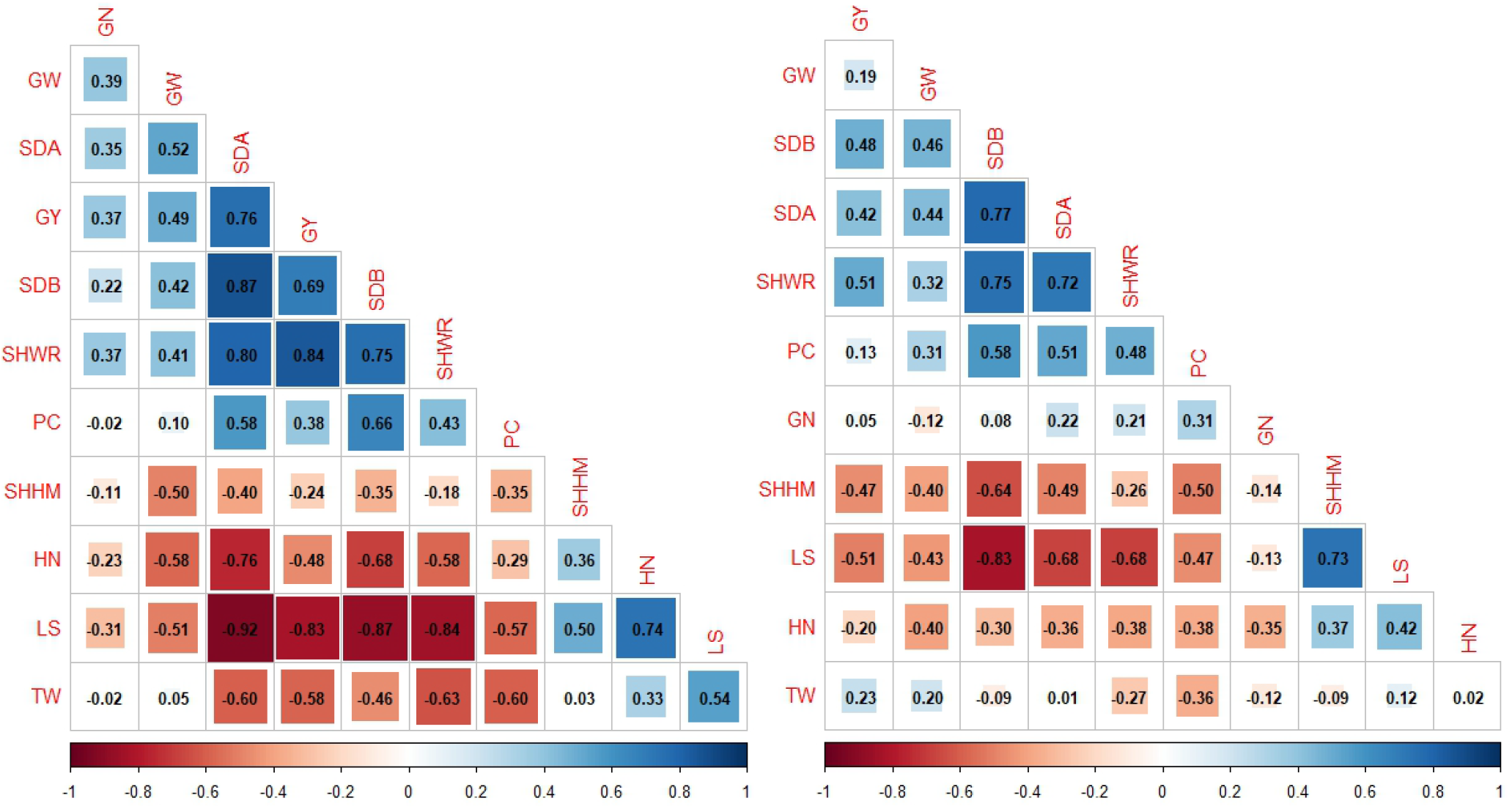
Combined correlation plots of Pearson’s correlation analysis among assessed attributes of soft white common spring wheat (SWCSW) variety Lousie under nitrogen fertilization (left) and under nitrogen in combination with PGR’s application (right). SHWR: stem height at water ripe; SHHM: stem height at the harvest maturity; SDA: stem diameter at water ripe (GS71); SDB: stem diameter at harvest maturity GS92; GN: grain numbers per head; HN: head numbers per fit; GY: grain yield; GW: 1000 grain weight; PC: protein contents; TW: test weight; LS: lodging score. Blue and orange shaded squares indicate positive or negative associations. Computed Pearson’s correlation coefficients are reported in the squares. The intensity of color shades and the size of colored squares are proportional to the computed Pearson’s coefficients.

The correlation among the studied attributes of Diva also varied under N fertilization and under application of PGR’s. Grain yield was positively correlated with stem height at water ripe (0.77), test weight (0.81), stem diameter at water ripe (0.67), grain weight (0.79) and stem diameter at harvest maturity (0.78) whereas it was negatively correlated with stem height at harvest maturity (-0.56) and lodging score (-0.83). Lodging score was positively correlated with grain weight (0.84), stem harvest at harvest maturity (0.85) whereas it was negatively correlated with stem height at water ripe (-0.67) test weight (-0.64), stem diameter at water ripe (-0.91), stem diameter at harvest maturity (-0.87), grain weight (-0.82) and protein contents (-0.55) (Figure 9). Considering the effect of PGR’s, association among the attributes of Diva was affected and intensity of correlation generally decreased in negative correlations and increased for positive correlations. Correlation between grain yield and lodging score (-0.73), stem diameter at water ripe (-0.67), stem diameter at harvest maturity (-0.79), stem height at water ripe (-0.49) generally decreased compared to under N fertilization. Lodging score and stem height at harvest maturity were positively correlated and intensity of correlation decreased with correlation coefficient of 0.71 under application of PGR’s compared to their positive association under N application with correlation coefficient 0.85. Stem height at harvest maturity negatively correlated with stem diameter at water ripe (-0.68) and stem diameter at harvest maturity (-0.81). A similar pattern of reduced negative association was also observed among other attributes (Figure 9).

**Figure 9.**
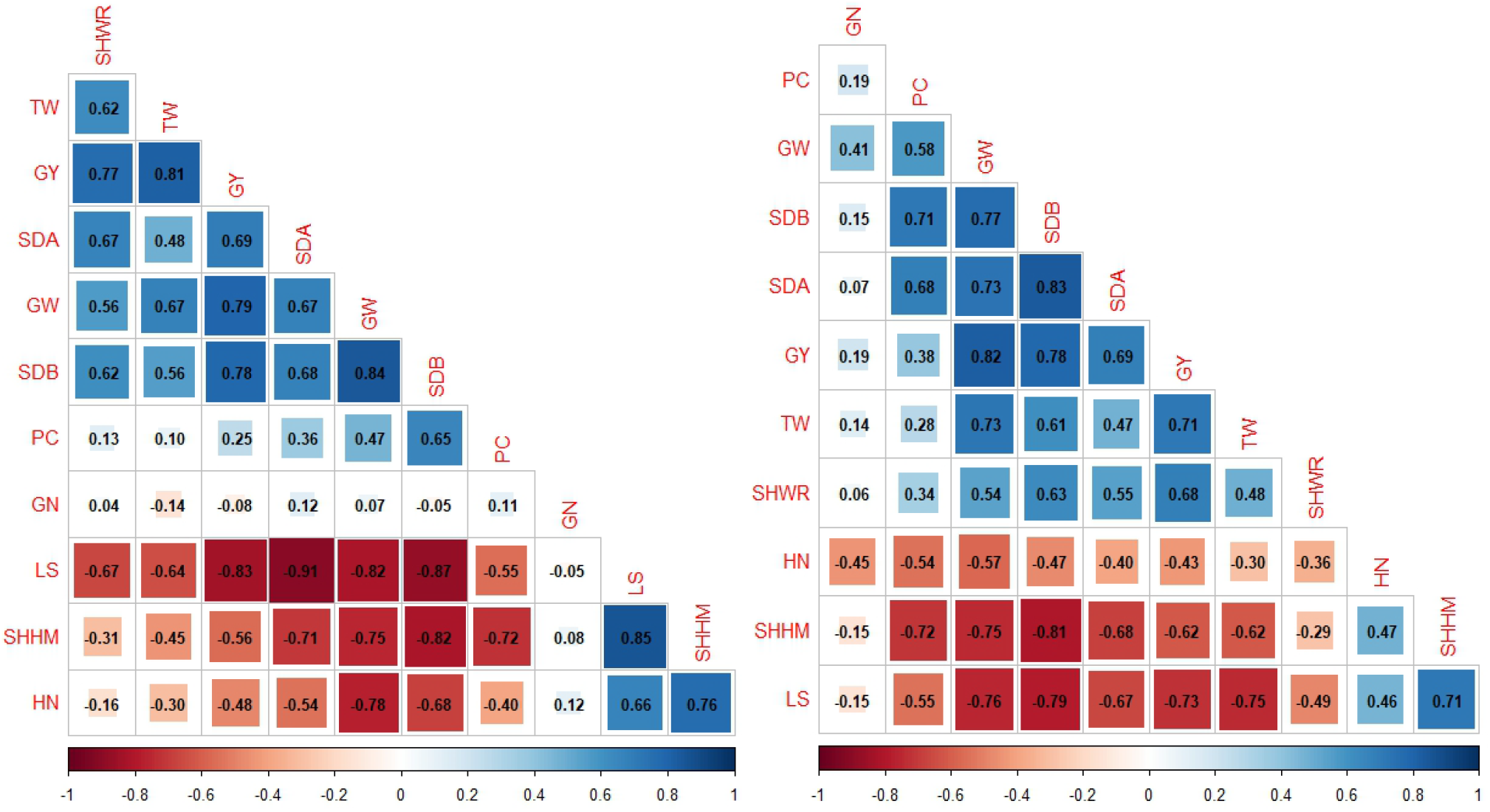
Combined correlation plots of Pearson’s correlation analysis among assessed attributes of soft white common spring wheat (SWCSW) variety Diva under nitrogen fertilization (left) and under nitrogen in combination with PGR’s application (right). SHWR: stem height at water ripe; SHHM: stem height at the harvest maturity; SDA: stem diameter at water ripe (GS71); SDB: stem diameter at harvest maturity GS92; GN: grain numbers per head; HN: head numbers per fit; GY: grain yield; GW: 1000 grain weight; PC: protein contents; TW: test weight; LS: lodging score. Blue and orange shaded squares indicate positive or negative associations. Computed Pearson’s correlation coefficients are reported in the squares. The intensity of color shades and the size of colored squares are proportional to the computed Pearson’s coefficients.

Correlation assessment indicated that under N fertilization, intensity of association among wheat attributes was higher than under application of PGR’s. This indicates that the interactive effect of wheat attributes was decreased on yield and quality parameters resulting in sustained yield concurrently affecting other attributes such as stem height which ultimately reduced logging.

## 5. Conclusions

The application of PGR’s resulted in a significant reduction in stem height of soft white common spring wheat (SWCSW) varieties. However, there was a little positive effect of PGRs on stem diameter at water ripe (SDA: GS71) and stem diameter at harvest maturity (SDB: GS92), grain numbers and head numbers. This effect indirectly contributed to the resistance of Louise and Diva against lodging, such as increased stem diameter enhanced stem strength. Although impact of PGRs application on grain numbers, head numbers grain yield and grain weight were non-significant and variable, we observed an improved performance of these attributes such as higher grain yield was observed under application of CC-B and CC-BC. In the first season no lodging was observed, whereas in the second season, we observed variable but positive response particularly under application of CC-C, TE, CC-BC and CC-AB. Lodging can be influenced by several factors such as stem thickness or diameter, stem height, grain weight, overall plant health and climatic conditions. We observed increased stem diameter under PGR application which indirectly contributed to stem strength and reduced lodging. Optimal nitrogen application is essential to avoid agronomic and economic losses. In this study we observed that the grain yield of Louise and Diva was similar at low and high N input. These results indicate that application of 168 kg N ha^-1^ is sufficient for an acceptable grain productivity and to gain agronomic and economic returns. We observed a significant seasonal impact and variations in the performance of SWCSW varieties, therefore, future research should consider long term evaluations to further get insights into the impacts of PGR’s on soft white common spring wheat varieties.

## Author Contributions

TH: Conceptualization (analysis and interpretation of the dataset), methodology, investigation, data curation, formal analysis, visualization, writing – original draft, writing – review and editing, project management.

## Acknowledgments

This work is part of one of the postdoctoral fellowship requirements and responsibilities at Oregon State University on publishing the existing dataset from Oregon State University that was provided to the author by the OSU Agronomy Program. Postdoctoral scholar Tajamul Hussain thanks Oregon State University for providing the resources to carry out this work.

